# bootRanges: Flexible generation of null sets of genomic ranges for hypothesis testing

**DOI:** 10.1101/2022.09.02.506382

**Authors:** Wancen Mu, Eric Davis, Stuart Lee, Mikhail Dozmorov, Douglas H. Phanstiel, Michael I. Love

## Abstract

*bootRanges* provides fast functions for generation of bootstrapped genomic ranges representing the null sets in enrichment analysis. We show that shuffling or permutation schemes may result in overly narrow test statistics null distributions, while creating new ranges sets with a block bootstrap preserves local genomic correlation structure and generates more reliable null distributions. It can also be used in more complex analyses, such as accessing correlations between cis-regulatory elements (CREs) and genes across cell types or providing optimized thresholds, e.g. log fold change (logFC) from differential analysis. The *bootRanges* functions are available in the R/Bioconductor package *nullranges* at https://bioconductor.org/packages/nullranges.

## 1 Introduction

In genomic analysis, to assess whether there is a significant association between two sets of ranges, one must choose an appropriate null model (De *et al*., 2014; Kanduri *et al*., 2018). For example, an enrichment of ATAC-seq peaks near certain genes may indicate a regulatory relationship (Lee *et al*., 2020), and enrichment of GWAS SNPs near tissue-specific ATAC-seq peaks may suggest mechanisms underlying the GWAS trait. Such analyses rely on specifying a null distribution, where one strategy is to uniformly shuffle one set of the genomic ranges in the genome, possibly considering a set of excluded regions. However, uniformly distributed null sets will not exhibit the clumping property common with genomic regions. Using an overly simplistic null distribution that doesn’t take into account local dependencies could result in misleading conclusions. More sophisticated methods exist, for example GAT, which allows for controlling by local GC content (Heger *et al*., 2013), and regioneR, which implements a circular shift to preserve clumping property(Gel *et al*., 2016). The block bootstrap (Politis *et al*., 1999) provides an alternative, where one instead generates random sets by sampling large blocks of ranges from the original set with replacement, as proposed for genomic ranges by Bickel *et al*. (2010) in Genome Structure Correlation (GSC). Using the block bootstrap is more computationally intensive than simple shuffling, and so GSC implements a strategy of swapping pairs of blocks to compute overlaps, while avoiding a genome-scale bootstrap.

Here we describe the *bootRanges* software, with efficient vectorized code for performing block bootstrap sampling of genomic ranges. *bootRanges* is part of a modular analysis workflow, where bootstrapped ranges can be analyzed at block or genome scale using tidy analysis with *plyranges* (Lee *et al*., 2019). We provide recommendations for genome segmentation and block length motivated by analysis of example datasets. We demonstrate how *bootRanges* can be incorporated into complex downstream analyses, including choosing the thresholds during enrichment analysis and single-cell multi-omics.

## 2 Features

*bootRanges* offers a simple “unsegmented” block bootstrap as well as a “segmented” block bootstrap: since the distribution of ranges in the genome exhibits multi-scale structure, we follow the logic of Bickel *et al*. (2010) and consider to perform block bootstrapping within *segments* of the genome, which are more homogeneous in their feature density. We consider various genome segmentation procedures or annotations, e.g. Giemsa bands or published segmentations (see Supplementary for details). The genome segments define large (e.g. on the order of ∼1 Mb), relatively homogeneous segments within which to sample blocks (Figure 1A). The input for the workflow is region sets ***x*** and ***y***, with optional metadata columns that can be used for computing a more complex test statistic than overlaps. Given a segmentation and block length *L*_*b*_, a *bootRanges* object is generated, which concatenates ranges across bootstrap iterations. This *bootRanges* object can be manipulated with *plyranges* to derive the bootstrap distribution of test statistics {*s*_*r*_} and a bootstrap p-value: 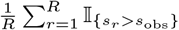 (Figure 1B). The *bootRanges* algorithms are explained schematically in Supplementary Section 1.6.

**Figure 1:**
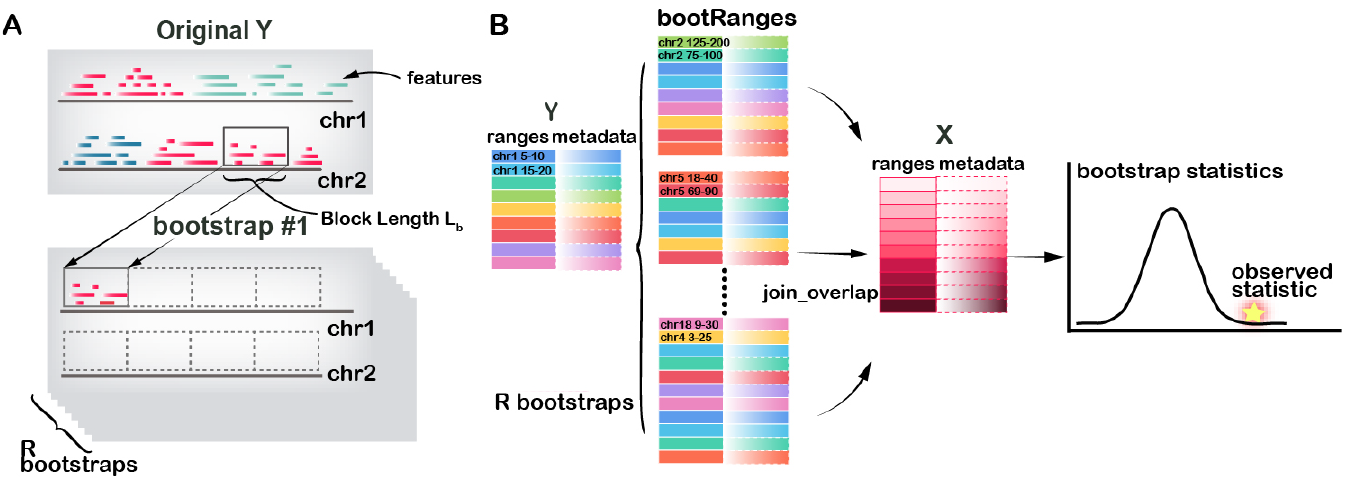
Overview of *bootRanges* workflow. A) Bootstrapping blocks of length *L*_*b*_. B) Testing overlaps of ranges in ***x*** with ranges in ***y***, comparing to a bootstrap distribution.

## 3 Application

We first applied *bootRanges* to determine the significance of the overlap of liver ATAC-seq (Currin *et al*., 2021) with SNPs associated with total cholesterol, bootstrapping the SNPs to assess significance. While the observed overlap was significant across many combinations of various segmentation methods and *L*_*b*_ according to empirical p-value, the variance of the bootstrap statistics distribution and the resulting *z* score varied greatly (Figure 2A-B). We used the *z* score to measure the distance between the observed value and bootstrap distribution in terms of standard deviations. Overlap rate was defined as the proportion of SNPs that had peaks within 10kb. That the variance of the distribution in Figure 2A for the unsegmented bootstrap increased with *L*_*b*_ indicated that ranges are inhomogeneous and bootstrapping with respect to a genome segmentation may be a more appropriate choice (Bickel *et al*., 2010). The decreasing trend using pre-defined segmentation from Roadmap Epigenomics indicated too many short segments, where *L*_*b*_ is too close to *L*_*s*_ for effective block randomization. To choose an optimal *L*_*b*_ and segmentation, we considered a number of diagnostics including those recommended previously (Bickel *et al*., 2010): the variance of the bootstrap distribution (Figure 2A), a scaled version of the changes in this variance over *L*_*b*_, and the inter-range distances distribution to assess conserved clustering of ranges (Supplementary Section 1.5). Here *L*_*s*_ ≈ 2 Mb and *L*_*b*_ ∈ [300kb, 600kb] was shown to be a good range for segment and block lengths (Figure S1A-C, and Supplementary Section 2.1). The scientific conclusion of this example was that liver ATAC-seq peaks were much closer to total cholesterol SNPs than expected even when placing blocks of SNPs to match a genome segmentation. Shuffling of genomic ranges (Supplementary Section 1.3) resulted in a much higher *z* = 18.5, compared to *z* ≈ 4 consistent with previous conclusions that shuffling may overestimate significance leading to misinterpretation of enrichment.

**Figure 2:**
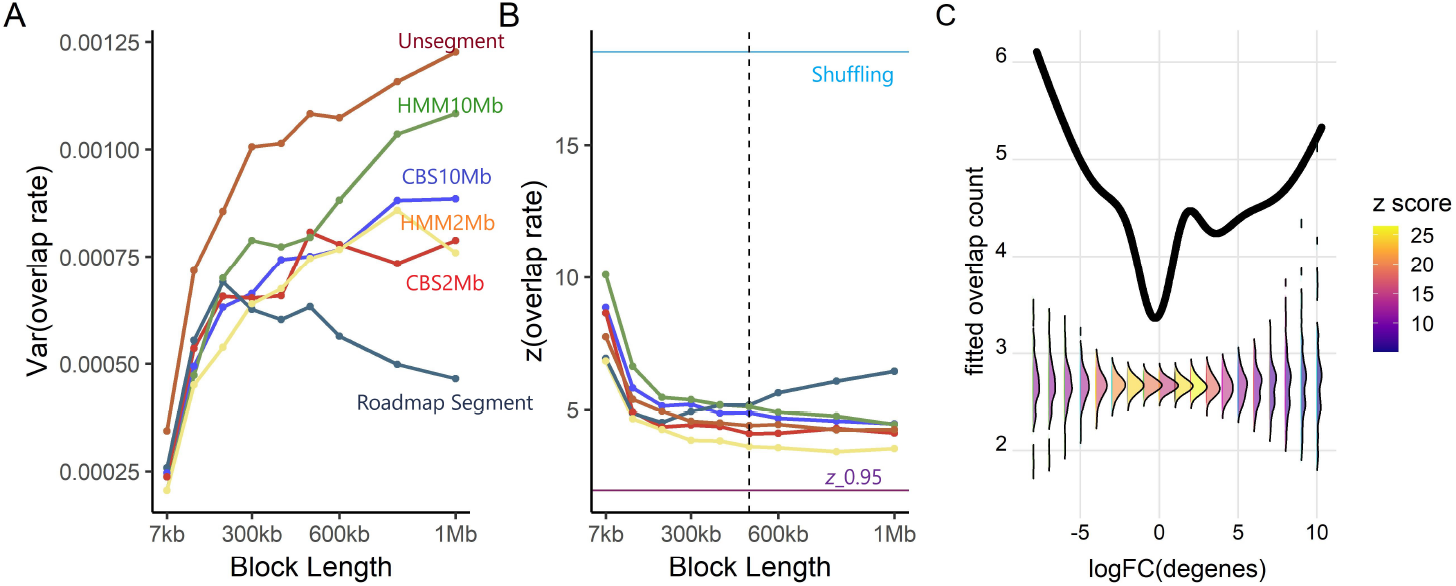
Parameter selection and overlap analysis. A) Variance of the rate of overlaps and, B) *z* score for the overlap, for different segmentations and *L*_*b*_ on the liver dataset. C) GAM predicted curves for observed (black line) and bootstrapped data (densities), for the overlap count over gene logFC. Conditional densities are colored by the *z* score for the overlap.

We demonstrated using *bootRanges* to motivate the choice of data-driven thresholds during enrichment analyses. We tested this on a dataset of differential chromatin accessibility and gene expression (Alasoo *et al*., 2018; Lee *et al*., 2020). A generalized linear model (GLM) with penalized splines was fit to the overlap count over gene logFC, both for the original data and to each of the generated null sets. Conditional densities of splines fit to null sets were computed at various thresholds to reveal how the threshold choice would affect the variance of the bootstrap density and the resulting *z* score (Figure 2C). These analyses suggested that |logFC| = 2 were optimized thresholds where the *z* score was highest (Figure S2B).

We applied *bootRanges* to Chromium Single Cell Multiome ATAC-seq and RNA-seq, to assess the correlation (*ρ*) of log counts for the two modalities for all pairs of genes and peaks, across 14 cell types (pseudo-bulk). Across all genes, we observed 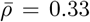, which was significantly higher than the bootstrap correlation mean (Figure S3A, 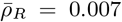). As expected, RNA and ATAC measured at local peaks had similar cell-type-specificity. Additionally, the average gene-peaks correlation per gene can be computed and compared to a bootstrap distribution to identify gene-promoter pairs that were significantly correlated across cell types (Figure S3B-C).

To compare speed, we ran *bootRanges* and GSC on ENCODE kidney and bladder ChIP-seq. The average time to block bootstrap the genome using *bootRanges* was 0.30s and 0.37s with overlap computation. A comparable analysis with GSC took 7.56s.

## 4 Conclusion

*bootRanges* efficiently generates null models of genomic ranges preserving local genomic correlations, and can be used easily in combination with other range-based tools such as *plyranges*. It has great flexibility in various disciplines (e.g. identify putative transcription factor binding site according to enriched peak regions).

## Data availability

All of the R code and data used in this paper are available at the following repository: https://github.com/Wancen/bootRangespaper. Companion website: https://nullranges.github.io/nullranges

## Acknowledgements

We thank Herve Pages for writting the prototype efficient vectorized code for bootstrapping, Haiyan Huang and Nancy Zhang for giving us insight into GSC methods, and Tim Triche for helpful feedback and suggestions.

