## Supplementary for "bootRanges: Flexible generation of null sets of genomic ranges for hypothesis testing"

### 1 Supplementary Methods

#### 1.1 Comparison to previous methods

In order to generate background ranges for hypothesis testing of association analysis, there are two general categories of methods. One strategy is to sample from a larger experimental pool or database. Methods following this strategy include LOLA (Sheffield and Bock, 2016) using Fisher's exact test, Poly-Enrich (Lee *et al.*, 2020a) using a likelihood ratio test based on a Negative Binomial likelihood, and matchRanges (Davis *et al.*, 2022) utilizing a covariate-based matching method. Another strategy is to permute or shuffle the genomic ranges, possibly considering an exclusion list of regions where the original region set should not be re-located. Example methods belonging to this category include bedtools shuffleBed (Quinlan and Hall, 2010), ChIP-Enrich (Welch *et al.*, 2014), and GenometriCorr (Favorov *et al.*, 2012). In addition, the method GAT allows controlling for GC content (Heger *et al.*, 2013), and regioneR implements a circular shift to preserve the clumping property of genomic ranges (Gel *et al.*, 2016). Our method falls in the second category in that we redistribute ranges along the genome, though we use a block resampling scheme to preserve local genome structure, as proposed for genomic region sets by Bickel *et al.* (2010).

#### 1.2 Segmentation

In the following, we define genome segmentation and how it pertains to block bootstrap resampling, as proposed by Bickel *et al.* (2010). The overall motivation for bootstrapping with respect to a segmented genome is to preserve large scale genomic structure, e.g. large regions with low or high density of genomic features such as genes. This preservation of structure through segmentation is in addition to the preservation of local clumping property of ranges, which is accomplished by resampling blocks.

In this work, we segmented the genome based on gene density. We downloaded the Ensembl v86 genes (Cunningham *et al.*, 2021) and counted the number of genes per million base pairs. We then supplied this vector of counts to various methods for segmentation. We sought to define sections of the genome that exhibit stationarity, where in this case, stationary means similar gene density within a segment. We additionally sought to group together segments across the genome that had similar gene density, such that a segmentation state $i \in [1, \ldots, S]$ consisted of one or more non-contiguous regions of the genome with similar gene density.

*

For block bootstrapping with respect to genome segmentations, we considered various genome segmentation methods, Circular Binary Search (CBS) (Olshen *et al.*, 2004) and a hidden Markov model (HMM) (Cardenas-Ovando *et al.*, 2017), as well as pre-defined segmentations, ChromHMM applied to Roadmap Epigenomics data (Ernst and Kellis, 2012). CBS and HMM have R package implementations, and so we incorporated these into a utility function in *nullRanges*, `segmentDensity()`. CBS, implemented in the *DNAcopy* package, was used to recursively split chromosomes into subsegments with similar density. In this case, the process was based on the square root of gene count per megabase to stabilize variance and reduce the heavy right tail of the distribution. We then used k-means to cluster chromosome subsegments into groups with similar gene density. We found that it was effective to cluster genome segmentations into three states, corresponding to low, medium, and high gene density. HMM was used to model the genes per megabase as an emission density with pre-defined number of states. We found that three provided effective segmentation. We then used the Viterbi algorithm for the hidden state decoding. Roadmap segmentations derived by ChromHMM (Ernst and Kellis, 2012) were downloaded from `https://egg2.wustl.edu/roadmap/web_portal/`. ChromHMM is based on a multivariate Hidden Markov Model that explicitly models the presence or absence of many chromatin marks. The original ChromHMM annotation generated segmentation including 15 small states. We then summarized these into 3 general categories: low density('E9','E13','E14','E15'), middle density('E10','E11','E12'), and high density('E1-E8'). After merging, our version of ChromHMM applied to Roadmap data had 8,797 ranges were left and the mean width was around 0.33 Mb.

#### 1.3 Simple region shuffling

Simple region shuffling was performed by placing ranges of interest uniformly in acceptance regions with probability proportional to the original SNP count per chromosome. For the paper, we defined acceptance regions as the inverse of ENCODE excluded regions (Amemiya *et al.*, 2019), centromeres, and telomeres from UCSC, as provided in the `excluderanges` Bioconductor package (Dozmorov *et al.*, 2022).

#### 1.4 Assessment of significance via bootstrap

Suppose we are interested in computing and assessing the significance of the overlap between one set of ranges $\boldsymbol{x}$ and another set of ranges $\boldsymbol{y}$. We compute the overlap $s_{obs}$, i.e. the total number of ranges in $\boldsymbol{x}$ that overlapped a range in $\boldsymbol{y}$. This statistic can vary in many ways, including computing the fraction of basepair overlap, restricting by a minimal amount of overlap, and allowing for a gap between the ranges, e.g. we may allow 1 kb of gap to connect local chromatin accessibility peaks to transcription start sites (TSS). Via block bootstrap resampling, we generate $R$ new sets of ranges $\boldsymbol{y_r}$ within $R_b = R \times \sum_{c=1}^{C} ceiling(L_c/L_b)$ blocks, for the chromosome $c$ length $L_c$. For details see Algorithm 1 and Algorithm 2. We can perform either genome-wise, block-wise, or region-wise analysis based on the question to be addressed. For example, if we want to test the significance of the genome-wide amount of overlap, we would compare to the genome-wise distribution. However, if we want to test the significance of the number of overlaps for a particular block or even a particular region, we could use a lower $R$ as there will be many more bootstrapped blocks and bootstrapped ranges for a block-wise or region-wise analysis. Without loss of generality, we will consider the genome-wise case. We use the overlaps per bootstrap sample $\boldsymbol{y_r}$ to calculate the overlap statistics $s_1, s_2, \ldots, s_R$ to derive a genome-wise empirical $p$-value $= \frac{1}{R}\sum_{r=1}^{R} \mathbb{I}_{\{s_r > s_{\text{obs}}\}}$ or $s_1, s_2, ..., s_{R_b}$ for block-wise analysis. Alternatively, we could use $z$ score to measure the distance between $s_{obs}$ and the set of $\{s_r\}$ in terms of standard deviations (SD). Specifically, the $z$ score in Figure 2 is calculated by

$$z = \frac{s_{obs} - \bar{s}}{\text{SD}_R},$$

where $\bar{s} = \frac{1}{R}\sum_{r=1}^{R} s_r$ is the sample mean of the overlap statistic $s_r$ over the $R$ bootstrap samples and $\text{SD}_R = \sqrt{\frac{1}{R-1}\sum_{r=1}^{R}[s_r - \bar{s}]^2}$ is the sample standard deviation. Note that, if block-wise analysis is preferred as in Bickel *et al.* (2010), we found out the SD of block-wise bootstrap distribution should be approximately scaled by $\sqrt{L_b}$ in simulation in order to make genome-wise and block-wise $z$ score comparative. In theory, the $\bar{s}$ is approximately the same in both analyses, but the SD is different. As $\text{SD}_R$ captures genome-wise SD, then block-wise SD is $\text{SD}_{R_b} = \sqrt{\frac{1}{R_b-1}\sum_{r=1}^{R_b}[s_r - \bar{s}]^2} \approx \sqrt{\frac{L_b}{L_G}} \times \sqrt{\frac{1}{R}\sum_{r=1}^{R_b}[s_r - \bar{s}]^2}$, for the whole genome length $L_G$.

#### 1.5 $L_b$ selection

We considered two aspects of the block bootstrap datasets to select block length $L_b$. First, following Bickel *et al.* (2010), we tried to find $L_b$ that provided the minimum value of a pseudo-metric,

$$d^*(v) = \left| \sqrt{\frac{L_{v-1}}{L_v}} [\mathrm{IQR}(\mathcal{L}_{L_v}) - \mathrm{IQR}(\mathcal{L}_{L_{v-1}})] \right|,$$

where $\mathcal{L}_{L_v}$ is the distribution of the test statistic at block length $L_v$, $v = 1, 2, \ldots, V$, $V$ is the number of candidate block lengths, and $\mathrm{IQR}(\mathcal{L})$ is the interquartile range of a distribution $\mathcal{L}$. Another aspect for block length selection is evaluating the conservation of the spatial pattern. In order to provide robust statistical inference in terms of significance testing, the generated null models should have similar spatial properties with the original sets, e.g. inter-range distances. Here we use "null models" interchangeably with block bootstrapped data. We used the Earth's Mover Distance (EMD) to quantify the similarity of the distributions of inter-range distances between the original and null models, resulting in values from zero (identical distributions) to one (totally disjoint distributions) (Rubner *et al.*, 1998). The EMD between two distributions is proportional to the minimum amount of "work" required to change one distribution ($y$) into the other ($y'$). Here $y$ and $y'$ are the histograms of original and null model inter-range distance. We observed that EMD always decreased as $L_b$ increased, because more neighboring ranges' distances were preserved in the block bootstrap with larger blocks (block boundaries break original distances). However, $L_b$ cannot be too large and close to $L_s$. Otherwise, the randomization of blocks would not be effective: the blocks will not sufficiently shuffle ranges across segments within a particular segmentation state. Hence, we decided an optimal $L_b$ falls in the range where EMD was around elbow of the line plot for this aspect.

#### 1.6 Block bootstrapping algorithm in *bootRanges*

**Algorithm 1:** Un-segmented block bootstrap / permutation

**Data:** Ranges ($\boldsymbol{y}$), Block length ($L_b$), Length of chromosome $c$ ($L_c$), Total number of chromosomes ($C$), Bootstrap iterations ($R$), Type (*permute* or *bootstrap*), Excluded regions ($\boldsymbol{e}$)

**Result:** `bootRanges` object, with the bootstrapped region sets from each iteration, concatenated across $R$ total iterations. Iteration and $L_b$ recorded as metadata

```
r ← 1
while r ≤ R do
    rearranged blocks ← Generate consecutive tiling blocks with width L_b (except last tile
      per chromosome which is cut to fit L_c for chromosome c)
    if permutation then
        random blocks ← Sample blocks without replacement from rearranged blocks
    else if bootstrap then
        n_b ← Σ_{c=1}^{C} ceiling(L_c/L_b)
        random blocks ← Sample n_b blocks with replacement from the genome, where
          sampling probability per chromosome is proportional to L_c
    bootstrap[r] ← Shift ranges in y that fall in the random blocks to their respective
      positions in the rearranged blocks, trimming applied to any ranges that extend past
      chromosome end or into excluded regions e
Concatenate bootstrap ranges across iterations r
```

**Algorithm 2:** Segmented block bootstrap with fixed or proportional block length

**Data:** Ranges ($\boldsymbol{y}$), Maximal block length ($L_b$), Length of genome ($L_G$), Bootstrap iterations ($R$), Type (*fixed* or *proportional*), Segmentation states $\mathcal{S}$, Segmentation regions $\mathcal{A}_s$ (the set of all ranges belonging to state $s$), and with $L_a^s$ representing the width of region $a$ in state $s$, Excluded regions ($\boldsymbol{e}$)

**Result:** `bootRanges` object

$r \leftarrow 1$
**while** $r \leq R$ **do**
  **foreach** *segmentation state* $s \in \mathcal{S}$ **do**
    $L_s \leftarrow \sum_{a \in \mathcal{A}_s} L_a^s$ ... thus $L_G = \sum_{s \in \mathcal{S}} L_s$
    **if** *fixed* **then**
      $n_b^s \leftarrow \sum_{a \in \mathcal{A}_s} \text{ceiling}(L_a^s / L_b)$
    **else if** *proportional* **then**
      $L_b^s \leftarrow L_b \cdot L_s / L_G$ ... thus $L_b^s \leq L_b$ and equal iff $\mathcal{S} \equiv \{s\}$
      $n_b^s \leftarrow \sum_{a \in \mathcal{A}_s} \text{ceiling}(L_a^s / L_b^s)$
    `rearranged blocks[s]` $\leftarrow$ Generate $n_b^s$ consecutive tiling blocks over $\mathcal{A}_s$
    `random blocks[s]` $\leftarrow$ Sample $n_b^s$ blocks with replacement from $\mathcal{A}_s$, where number of blocks per $a$ is proportional to $L_a^s$
  Concatenate `rearranged blocks` and `random blocks` across $s$
  `bootstrap[r]` $\leftarrow$ Shift ranges in $\boldsymbol{y}$ as in Algorithm 1
Concatenate `bootstrap` ranges across iterations $r$

### 2 Supplementary Results

#### 2.1 Overlap analysis of liver ATAC-seq peaks with GWAS SNPs

1,872 SNPs were download from the NHGRI-EBI GWAS catalog (Buniello *et al.*, 2018) on September 22, 2021. We only extracted single nucleotide variants associated with "total cholesterol". Liver ATAC-seq peaks (Currin *et al.*, 2021) aggregated from 20 samples were downloaded from the GEO accession: GSE164870. All the peaks here were filtered by FDR adjusted $p$-value $\leq 0.05$. Then, genomic coordinates of consensus peaks were converted from hg19 to hg38 using `liftOver` to obtain 221,606 peaks.

Based on the $L_b$ selection criteria, we found that $d^*$ had smaller values in most cases when $L_b \in [300kb, 800kb]$ (Figure S1A). When quantify the similarity of the distributions of inter-range distances between the original and null models using EMD, we derived the histograms of original and null model inter-range distance with bin size = 0.3 under various $L_b$ and $L_s$ settings. For this aspect, $L_b \in [200kb, 600kb]$ was shown to be a good range by visualization of density plot (Figure S1B) and according to the elbow of the EMD statistic (Figure S1C). Combining two aspects, we concluded that $[300kb, 600kb]$ was a good range for block length.

#### 2.2 Enrichment analysis of macrophage ATAC-seq and gene expression

We used the differential results for RNA-seq and ATAC-seq as described in the *fluentGenomics* workflow (Lee *et al.*, 2020b), with the original data from a human macrophage experiment by Alasoo *et al.* (2018). The contrast of interest was interferon gamma (IFNg) treatment. Since the transcriptomic response to IFNg stimulation may involve increased transcription factor binding to *cis*-regulatory elements near genes, and as ATAC-seq captures the accessibility and activity at these regions, Lee *et al.* (2020b) hypothesized and demonstrated an enrichment of differentially accessible (DA) ATAC-seq peaks near differentially expressed (DE) genes. While Lee *et al.* (2020b) assessed significance by randomly sub-setting the non-DE genes and re-computing overlap with DA peaks, here we considered the block bootstrap for statistical significance.

When performing block bootstrap on the number of overlaps, using $L_b = 500kb$ and $R = 100$, we obtained empirical $p$-value = 0 and $z = -108.1$. We therefore rejected the null hypothesis and concluded that there was significant enrichment of DA peaks near DE genes.

For fitting the generalized penalized splines, we used the `gam` function in the `mgcv` package (Wood, 2011) to fit the model, based on a penalized likelihood maximization. Generalized cross-validation was used to choose the optimal value for the smoothing parameter, $\lambda$. Then, the `tidymv` package was used to predict and extract the fitted value. For each generated null set, we found all the predicted overlap counts for genes with logFC falling in the range (-8,10) with 1 as the increment. Then, conditional density plots can be plotted across all null sets. Additionally, we used a $z$ score to measure the distance of predicted overlap counts between observed sets and the null sets in terms of SD of the conditional null set (conditioning on logFC bin).

The following code shows how the analysis of Fig 2C is performed as the downstream task following *bootRanges*. Suppose $x$ contains the DE genes and *bootRanges* contains the bootstrapped DA peaks. For each gene within each iteration, we extract the total number of peaks that overlap with that gene and that gene's logFC (below, `max` is used to reduce a set of repeated, identical values to a single numeric). Then, after fitting splines, we predict overlap counts for a 2,000 point grid spanning the original logFC range. These spline-predicted values are those used to define the conditional density.

```
boot_stats <- x %>% plyranges::join_overlap_inner(bootRanges) %>%
  plyranges::group_by(id.x, iter) %>%
  plyranges::summarize(count = plyranges::n(), logFC = max(x_logFC)) %>%
  as.data.frame() %>%
  tidyr::complete(id.x, iter, fill=list(count = 0)) %>%
  dplyr::select(iter, count, logFC) %>%
  tidyr::nest(-iter) %>%
  dplyr::mutate(
     fit= map(data, ~gam(count ~ s(logFC), data = ., family=poisson)),
         pred  = map(fit, ~predict_gam(model = ., length_out = 2000)),
         fitted = map(pred, ~find_fit(data=., logFC = seq(-8,10,1))))
```

#### 2.3 Correlation analysis of Single Cell ATAC + Gene Expression

We were interested in demonstrating *bootRanges* on data from single cell experiments, in particular looking at correlations at pseudo-bulk resolution in single cell multi-omics datasets. For this purpose, data were downloaded from Ricard Argelaguet and Marioni (2020), including RNA-seq for genes and ATAC-seq for peaks in 10,032 cells. Cell type annotations had already bee performed by the 10x Genomics R&D team. Information on genes and peaks from chromosomes 1-22 were selected. First, we aggregated data from cells within same cell type, to form pseudo-bulks data with $n = 14$ cell types according to the metadata. The pseudo-bulk data provided smoother correlation statistics without loss of the information of interest (correlation of cell-type-specific expression and accessibility). Next, we removed all the ranges with 0 standard deviation in either assay. Finally, log counts per million (CPM) were computed from `edgeR` (Robinson *et al.*, 2010) to account for different library sizes in both assays.

Our expectation was that the two modalities would have significantly high correlation for peaks close to genes ($le1kb$). For the whole gene set, the mean correlation of gene RNA-seq and peak ATAC-seq read counts was 0.33, while the bootstrap sampling correlation distribution in Figure S3A had a mean correlation of 0.007 for $R = 1,000$. Therefore, as expected, RNA and ATAC measured at nearby peaks had similar cell-type-specificity. Additionally, the average gene-peaks correlation per gene can be computed and compared to a bootstrap distribution to identify 5,644 gene-promoter pairs that were significantly correlated across cell types. Among these, 5,591 genes had only one paired promoter, while 25 genes had 2 nearby promoters (peaks of accessibility within 1kb of the body of a gene). Two examples of the most significant gene-peak pairs are shown in Figure S3B-C.

The following code is an example of how such an analysis can be performed with *bootRanges* and *plyranges*. Here $x$ refers to RNA-seq and $y$ refers to ATAC-seq.

```
# split count matrix into NumericList metadata and sort
x <- x_Granges %>%
  mutate(counts_X = NumericList(asplit(x.cpm, 1))) %>% sort()
y <- y_Granges %>%
  mutate(counts_y = NumericList(asplit(y.cpm, 1))) %>% sort()

# standardize read counts for fast correlation computation
x$counts_x <- NumericList(lapply(x$counts_X, function(z) (z-mean(z))/sd(z)))
y$counts_y <- NumericList(lapply(y$counts_y, function(z) (z-mean(z))/sd(z)))

# generate block bootstrap ranges for 'y'
boots <- bootRanges(y, blockLength = 5e5, R = 100)

# for standardized x and y:
correlation <- function(x,y) 1/(length(x)-1) * sum(x*y)

# extract bootstrap summary statistics
boot_stats <- x %>% plyranges::join_overlap_inner(boots, maxgap=1000) %>%
  plyranges::mutate(rho = correlation(counts_x, counts_y)) %>%
  plyranges::group_by(iter) %>%
  plyranges::summarise(meanCor = mean(rho))
```

### 3 Supplementary Figures

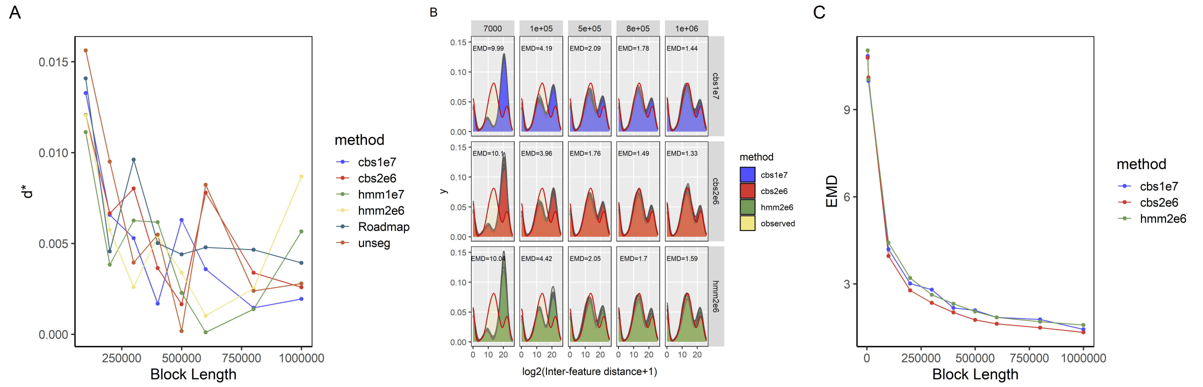


Figure S1: $L_b$ selection assessment. A) A pseudo-metric $d^*$ (Bickel *et al.*, 2010) over $L_b$ , B) log2(inter-range distance+1) density plots over various $L_b$ and segmentation settings. Red curve represents the observed ranges' log2(inter-range distance+1) density. The more a particular null set's density plot overlapped with observed ranges, the better we conserved the spatial distribution of the original set via block bootstrapping. Median EMD shown as text in each panel. C) Median EMD over $L_b$ where EMD quantified the similarity of inter-range distance distributions between nulls sets and observed sets.

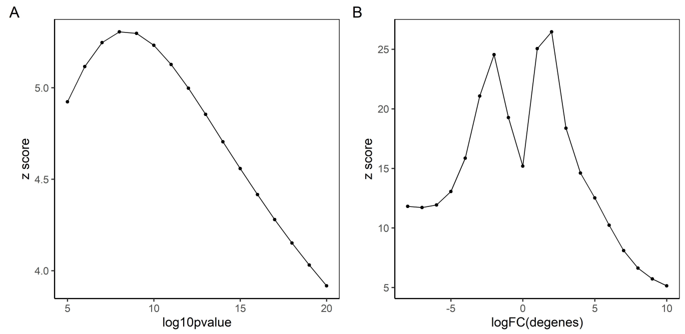


Figure S2: $z$ scores against thresholds for liver ATAC-seq and macrophage experiments. A) $z$ score against SNPs -log10 ($p$-value) in the liver dataset. The penalized splines was fitted on the top 1800 SNPs and was used to predict the overlap rate for the -log10 ($p$-value) falling into the range (5,20) with 1 as increment for both observed and null sets data. Then, $z$ scores, indicating the distance of the predicted overlap rate between the observed set and null sets($R = 1,000$) in terms of SD, was derived. B) $z$ score against gene expression logFC of the macrophage dataset. The penalized splines was fitted on all DE genes and was used to predict the overlap count for the logFC falling into the range (-8,10) with 1 as increment for both observed and null sets data. Then, $z$ scores, indicating the distance of the predicted overlap count between the observed set and null sets($R = 1,000$) in terms of SD, was derived.

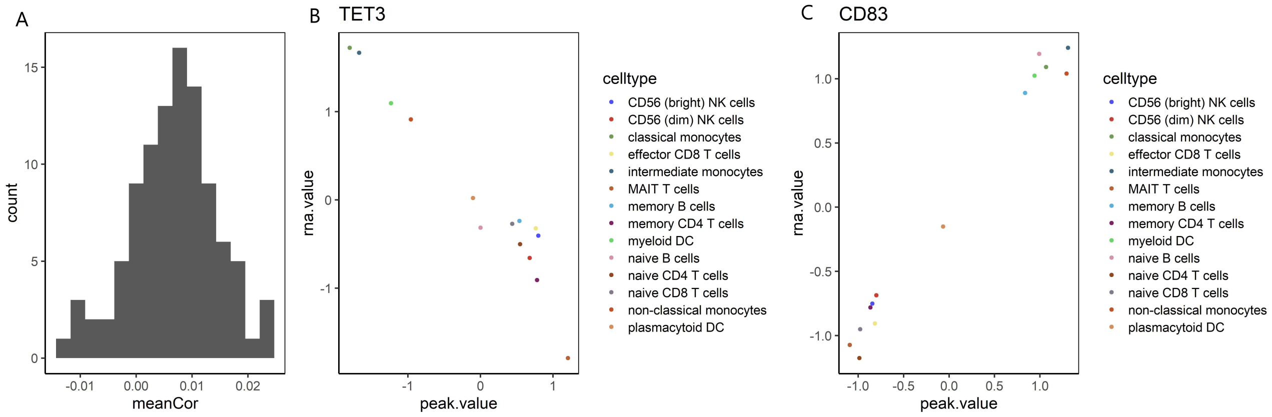


Figure S3: Correlation analysis of Chromium Single Cell Multiome ATAC + Gene Expression dataset. A) The distribution of mean correlation of gene expression with bootstrapped ATAC-seq peaks and their read counts ($R = 100$). B) TET3 read counts ("rna value") over peak chr2:74000098-74003475 read counts ("peak value"), colored by cell type. TET3 had the most negative observed correlation $\rho = -0.963$. C) CD83 read counts over peak chr6:14116971-14139988 read counts, colored by cell type. The correlation of this gene-promoter pair was 0.992 and was the highest among all the pairs.
